# New Insights into the Expression Landscape of the Genetic Hearing Loss Protein Whirlin Across Human Tissues and Cell Types

**DOI:** 10.64898/2026.03.23.713511

**Authors:** Aikaterini Dandini, Loren Méar, Cecilia Lindskog, Candice Gautier

**Affiliations:** Department of Chemistry for Life Sciences, Uppsala University, Uppsala 75123, Sweden; Department of Immunology, Genetics and Pathology, Cancer Precision Medicine Research Unit, Uppsala University, SE-75185 Uppsala, Sweden; Department of Women’s and Children’s Health, Division for Neonatology, Obstetrics, Gynaecology and Reproductive Health, Karolinska Institute and Karolinska University Hospital, SE-14186 Stockholm, Sweden; Centre of Excellence for Chemical Mechanisms of Life, Department of Chemistry for Life Sciences, Uppsala University, Uppsala 75123, Sweden

**Keywords:** Whirlin, DFNB31/WHRN, Usher syndrome, Genetic hearing loss, Human Protein Atlas, tissue expression, immunohistochemistry, transcriptomics, ciliated cells, endocrine cells

## Abstract

Whirlin (*WHRN*) is a scaffolding protein with essential roles in the development and function of specialised cells involved in hearing and vision. Although previous studies have primarily focused on the inner ear and retina, whirlin’s distribution across human tissues and its potential roles beyond these sensory systems remain poorly defined. Here, we present a comprehensive expression map of whirlin by integrating bulk RNA, single-cell RNA, and antibody-based proteomics data generated as part of the Human Protein Atlas consortium. *WHRN* was detected in most human tissues, with the highest levels observed in endocrine organs, reproductive tissues, and the nervous system. Single-cell analysis revealed that this distribution is driven by enrichment in specific cell types, including endocrine, ciliated, and specialised epithelial cells. Genome-wide specificity analysis placed *WHRN* amongst the most cell-type-specific genes in both ciliated and endocrine cell groups, and several of whirlin’s known binding partners also showed cell-type specificity in these groups. Together, these findings substantially broaden the known expression landscape of whirlin beyond established sensory systems and identify a previously unexplored association between Usher genes and endocrine cells.

## INTRODUCTION

Whirlin is a scaffolding protein originally characterised for its essential roles in the development and function of sensory systems, particularly hearing, balance, and vision.[1,2] It was initially identified in studies of whirler mice presenting profound hearing loss and vestibular dysfunction, and associated with genetic alterations on mouse chromosome 4 in the gene later named *Dfnb31/Whrn*.[3] In humans, variants in *WHRN/DFNB31* on chromosome 9q32-q34 are associated with autosomal recessive non-syndromic hearing loss, [3] and Usher syndrome type 2 (USH2), which is the most common cause of combined hereditary deafness and blindness.[4]

Whirlin comprises multiple protein-protein interaction domains including PDZ domains and Harmonin Homology domains (HHD) both in the N- and C-terminal region, and exists in distinct isoforms generated by alternative splicing. In the inner ear, it is part of the protein network that shapes the stereocilia bundles of inner and outer hair cells in the organ of Corti, the sensory epithelium responsible for hearing. Whirlin localises to two key subcellular regions of these bundles, including the tip, where it participates in the elongation and maintenance of the stereocilia, and the base, where it forms the USH2 complex with other USH2 proteins ADGRV1 (gene *ADGRV1*), Usherin (gene *USH2A*), and PDZD7 (gene *PDZD7*) crucial for bundle organisation during development. Vestibular hair cells have similar bundles where whirlin also regulates stereocilia elongation and maintenance.[5] Pathological variations in *WHRN* cause shortened, disorganised and consequently dysfunctional stereocilia observed in whirler mouse models,[6,7] consistent with its role in genetic hearing loss and balance dysfunction. In the retina, whirlin localises at the periciliary membrane of photoreceptors where it participates in the recruitment of other proteins to form the periciliary membrane complex (PMC), thought to support cargo transport to the connecting cilium.[7]

Although whirlin primarily received attention for its role in sensory cells, its expression in non-sensory tissues is not a recent discovery. In the same year that Mburu et al. identified whirlin as the protein essential for stereocilia elongation in the whirler mouse and responsible for DFNB31-related hearing loss, Yap et al. independently isolated the same protein in the brain, naming it CIP98, and described it as a synaptic scaffolding PDZ protein widely expressed in the central nervous system.[8] Since then, evidence has emerged supporting its implication in neuronal signalling, structure, and stability,[9] as well in proprioception[10] and nociception.[11] In addition, *WHRN/DFNB31* has appeared in transcriptomic and genetic studies across a range of clinical reports, including bipolar disorder,[12,13] cardiac dysfunction,[14] Hashimoto’s thyroiditis,[15] diabetes,[16] ciliopathies involving kidney and liver function,[17] and colorectal cancer.[18] These observations collectively suggest that whirlin may contribute to the function of cells other than those specifically involved in hearing, balance, and vision, yet investigations into its potential role in other diseases remain largely overlooked.

The Human Protein Atlas (HPA) is a large-scale, publicly accessible resource that maps the spatial distribution of human RNA and proteins across organs, cell types, and subcellular compartments using transcriptomics data combined with antibody-based proteomics. For several Usher syndrome proteins, including whirlin, protein expression data across different human tissues have so far been limited. Here, we present the first detailed body-wide expression analysis of whirlin, integrating bulk RNA, single-cell RNA, and antibody-based protein expression data from normal and cancerous tissues.[19–21] The underlying data was made available in the public version of the database (version 25). Single-cell RNA analysis revealed pronounced enrichment in both ciliated and endocrine cell types, and genome-wide specificity analysis placed *WHRN* among the most cell-type-specific genes in both groups. We further examined whether known sensory binding partners of whirlin showed similar cell-type specificity. Most notably, *ADGRV1* and *USH2A*, the known partners forming the USH2 complex with whirlin, displayed a comparable ciliary and/or endocrine specificity. Finally, we identified cell lines expressing endogenous *WHRN*, providing potential cellular models for investigating whirlin function and supporting future development of *WHRN*-targeted gene therapies. Such models may help address the lack of established cell lines for inner ear hair cells and photoreceptors, which is a longstanding limitation in the field. This work establishes a new foundation for investigating whirlin’s broader physiological role and potential pathological implications beyond sensory systems.

## RESULTS

### Whirlin expression extends well beyond sensory tissues at both transcript and protein levels

To evaluate the distribution of *WHRN* across the human body, we analysed transcriptomics from two complementary sequenced mRNA libraries (bulk RNA) retrieved from the HPA database covering all major systems including 51 tissues across 15 organ systems: the internally generated HPA dataset[20] and the Genotype-Tissue Expression (GTEx) dataset.[22] Figure 1a shows an overview of *WHRN* bulk RNA expression levels across all human tissues based on HPA and GTEx datasets.

**Figure 1:**
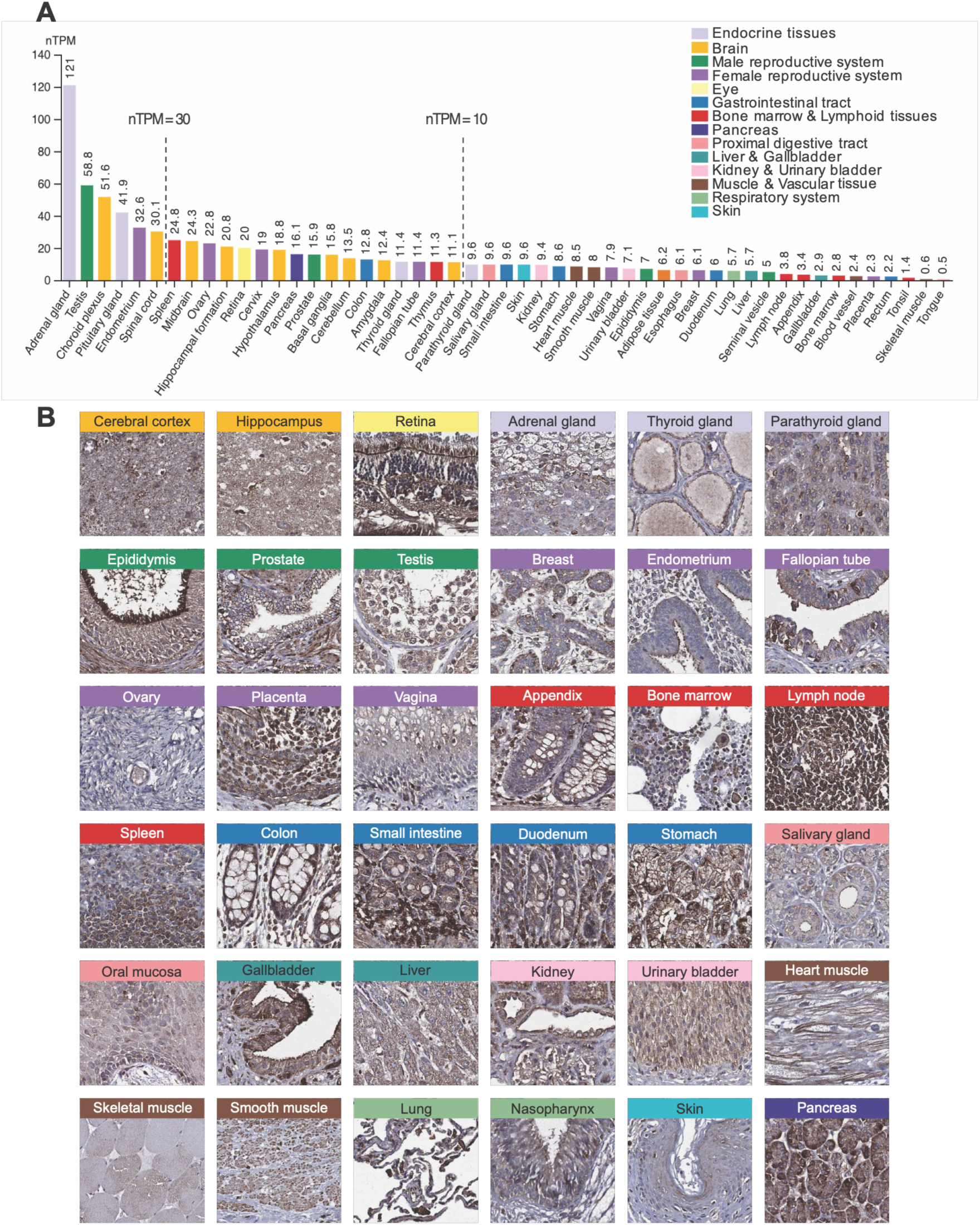
*WHRN* bulk RNA and protein expression across normal human tissues. (a) Bar plot showing the consensus normalised transcript per million (nTPM) values of *WHRN* bulk RNA across 51 human tissues ordered by decreasing expression level. (b) Representative IHC images per tissue type showing protein expression. Manually annotated protein expression levels per tissue are provided in Supplementary Table 1b.

Across the combined datasets, *WHRN* was detected in 49 out of 51 tissues, indicating a near-ubiquitous expression pattern. Among all tissues examined, the highest expression levels (>30 nTPM) were observed in endocrine tissues, reproductive tissues, and the brain and nervous system. Consistent with its established role in vision, the retina also exhibited relatively high *WHRN* abundance (20 nTPM). Beyond the major sites of expression, moderate *WHRN* levels (>10 nTPM) were observed in the spleen, pancreas, prostate, colon and thymus, whereas most tissues with squamous epithelial, as well as muscles and vascular tissues, consistently displayed low expression levels. Inner ear tissues were not represented in the datasets used in this study and therefore could not be assessed in this analysis.

To determine whether *WHRN* bulk RNA levels translated into detectable protein expression, we analysed whirlin expression by immunohistochemical tissue staining (IHC) in a panel of 45 normal human tissues, as well as extended samples of human retina. The data was generated using the stringent antibody validation workflow of the HPA, and made publicly available in the database version 25. Figure 1b presents an overview of representative IHC stained images. Consistent with transcriptomic analyses, whirlin was detected in multiple organ systems and displayed a broad distribution across different tissues. Strong staining was observed in the retina, particularly within photoreceptors, providing further support for the specificity of the antibody used in this study.

Whirlin was noticeably present in most tissues containing ciliated epithelia, including the brain, respiratory tract and multiple reproductive tissues including the endometrium, fallopian tubes and particularly strong staining in the stereocilia of the epididymis. Beyond these tissues, whirlin was also detected in a variety of glandular epithelial tissues, including endocrine, colon, stomach, duodenum, salivary gland, pancreas, and kidney tissues. In these organs, staining was primarily observed in epithelial cells lining the glandular and tubular structures rather than being uniform throughout the tissue. Interestingly, several cell types showed higher expression in the luminal membrane, including those from the placenta, salivary ducts, and gall bladder epithelium, as well as glandular cells of the gastro-intestinal tract. Whirlin was further detected in several immune-related tissues, including lymph node, spleen, and bone marrow, as well as in immune cells present throughout many other tissues.

### Single-cell RNA analysis reveals cell type enrichment of *WHRN* in ciliated and endocrine cells

To refine the tissue expression patterns observed in bulk RNA analyses and IHC staining, we examined the single-cell RNA sequencing data (scRNA) from the HPA database encompassing 34 different datasets and providing *WHRN* expression profiles across 154 distinct cell types.[19] In contrast to the bulk RNA analysis, which provides an average expression level for each tissue, the single-cell dataset allowed us to resolve the specific cell types in which the expression is most prominent within individual tissues.

Overall, *WHRN* was not uniformly expressed across different cell types and instead showed clear enrichment in specific cells. Figure 2a displays *WHRN* scRNA expression levels in the 26 cell types exceeding the 50 nCPM threshold. A complete list of nCPM values for all cell types is provided in Supplementary Table 2.

**Figure 2:**
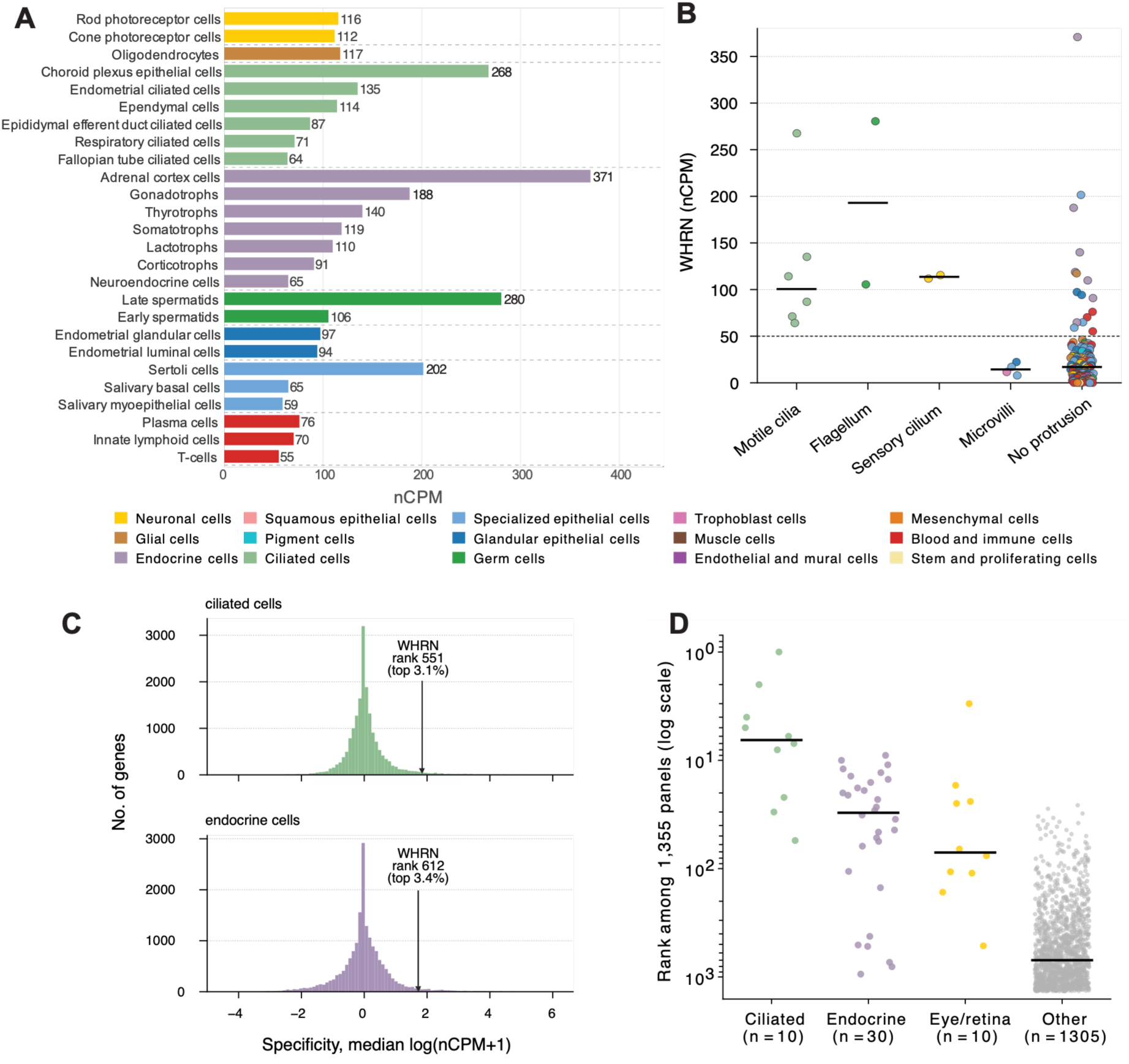
*WHRN* RNA expression at single cell level. (a) scRNA expression in human single cell types with nCPM>50. Normalised count per million (nCPM) values from the HPA single cell resource. Cell types are grouped based on similarities in histology and function. The entire dataset can be found in Supplementary Table 2.(b) Dot plot of scRNA levels (nCPM) across the 154 cell types, grouped by the type of protrusion each cell type is known to bear (motile cilia, flagellum, sensory cilium, microvilli, or no protrusion); dots are coloured by the same 15 cell type groups as in (A). Horizontal bars indicate the group median. The dashed line marks the 50 nCPM threshold. (c) Histograms showing the gene specificity of *WHRN* in ciliated and endocrine cells types. (d) Dot plot showing the rank (log scale; rank 1 = strongest enrichment) of each of the 1,355 WebCSEA reference panels by the combined-P enrichment of the top 500-gene co-expression module for *WHRN*. Panels are grouped into ciliated (n = 10), endocrine (n = 30), retinal (n = 10), and all remaining panels (n = 1,305). Horizontal bars indicate the group median rank.

The highest levels were observed in endocrine cells, with adrenal cortex cells displaying the strongest expression in the entire single-cell dataset (371 nCPM). Elevated expression was also observed in several hormone-producing pituitary cell types, including gonadotrophs (188 nCPM), thyrotrophs (140 nCPM), somatotrophs (119 nCPM), lactotrophs (110 nCPM), and corticotrophs (91 nCPM). Together, these findings are consistent with the high bulk RNA levels observed in endocrine tissues. Moderate scRNA levels were further observed in neuroendocrine cells (65 nCPM).

A second prominent enrichment was detected in ciliated cells and, more broadly, cells bearing specialised surface protrusions (Figure 2b). All cell types with motile cilia showed scRNA levels above 50 nCPM, and the expression was significantly higher in this cell type group than in the remaining cell types (Mann-Whitney U test, FDR = 0.0035; Fisher’s exact test for the proportion exceeding 50 nCPM, FDR = 0.0008). This accounts for the presence of *WHRN* in tissues containing specialised epithelia such as the brain (choroid plexus epithelial cells, 268 nCPM and ependymal cells, 114 nCPM), reproductive organs (endometrial ciliated cells, 135 nCPM; epididymal efferent duct ciliated cells, 87 nCPM; fallopian tube ciliated cells, 64 nCPM) and respiratory tissues (respiratory ciliated cells, 71 nCPM). Retinal rod and cone photoreceptors, which are neuronal cells containing specialised sensory cilia, also demonstrated elevated expression (116 and 112 nCPM respectively), supporting the established role of whirlin in vision. Male reproductive specialised cells, including both germ cells (early spermatids, 106 nCPM; and late spermatids, 280 nCPM), as well as somatic Sertoli cells (202 nCPM), showed markedly increased scRNA levels compared to most other cell types, consistent with the strong bulk RNA expression observed in male tissues. Immune cells, including plasma cells and lymphocytes, also showed moderately high expression levels (respectively 76 and 70 nCPM, respectively), likely contributing to the intermediate bulk RNA levels observed in lymphoid tissues such as spleen and thymus.

Interestingly, *WHRN* ranked amongst the most cell-type-specific genes in both ciliated and endocrine cells. A one-sided Mann-Whitney U test performed for each gene placed *WHRN* at rank 551 of 17,984 genes in ciliated cells, and rank 612 of 17,984 genes in endocrine cells (top 3.1% and 3.4%; FDR = 8.8 × 10⁻⁵ and 6.8 × 10⁻³, respectively) (Figure 2c). Only 0.56% of the 17,984 genes were at least as specific as *WHRN* for both cell families.

To further investigate the cell-type associations of *WHRN,* we performed a cell-type-specific enrichment analysis (CSEA). A 500-gene co-expression module was first defined, encompassing the 500 genes most strongly correlated with *WHRN* across the 154 cell types (Spearman ρ ≥ 0.37). This module showed strong enrichment for cilium-related Gene Ontology annotations and for Reactome pathways, including cilium assembly, intraflagellar transport, basal-body anchoring and BBSome-mediated cargo targeting. The 500-gene module was subsequently submitted to WebCSEA (https://bioinfo.uth.edu/webcsea/),[23] comprising 1,355 independent cell-type reference panels derived from published single-cell atlases (Figure 2d). Ciliated cell panels ranked the highest across the entire compendium (n = 10; median rank 6.5 of 1,355; Mann-Whitney P = 2.7 × 10⁻^8^), followed by endocrine cell panels (n = 30; median rank 30.5; Mann-Whitney P = 3.3 × 10⁻¹^4^); and retinal cell panels (n = 10; median rank 71; Mann-Whitney P = 8.4 × 10⁻⁷). Together, these three families comprised only 50 of the 1,355 panels (3.7%), but accounted for 29 of the 33 panels that passed the Bonferroni-adjusted significance threshold (WebCSEA combined P < 3.7 × 10⁻⁵).

### Whirlin’s principal binding partners share ciliated and endocrine cell-type specificity

To determine whether whirlin’s known binding partners exhibit similar cell-type expression specificity, we examined the ciliary and endocrine specificity rankings of each partner among the same 17,984 genes used to rank *WHRN* (Figure 3a). *ADGRV1*, encoding whirlin’s binding partner within the USH2 complex, displayed a dual specificity pattern resembling that of *WHRN*, ranking amongst the most cell-type-specific genes genome-wide for both ciliated (rank 427, FDR = 0.025) and endocrine (rank 97, FDR = 0.015) cells. This concordance was not shared by the complex’s other core component, *USH2A*, which reached significance for endocrine specificity alone (rank 815, FDR = 0.016; ciliary FDR = 0.070). Two other binding partners showed significant specificity for a single family, including *PCDH15* for endocrine cells only (rank 185, FDR = 0.019), and *ESPN* for ciliated cells only (rank 317, FDR = 0.016). None of the remaining examined partners reached statistical significance for either cell family.

**Figure 3:**
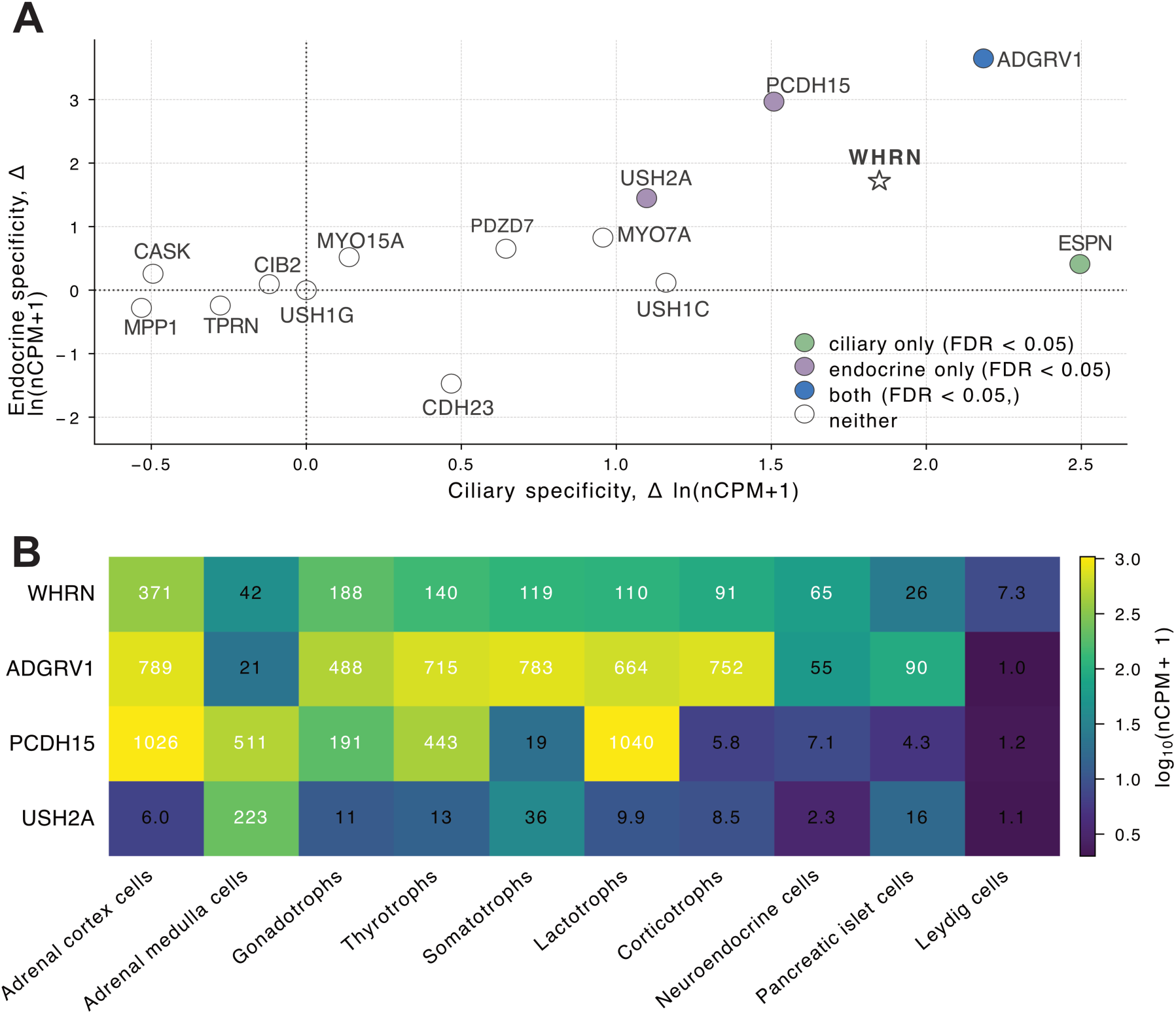
Whirlin binding partners and their cell-type RNA expression specificity. (a) Scatter plot of cell-type RNA expression specificity (Δ median ln(nCPM+1), family − rest of atlas) for 13 known binding partners. Ciliary specificity is shown on the x-axis and endocrine specificity on the y-axis. Partners are coloured according to significance (FDR < 0.05) in each family: ciliary-only, endocrine-only, both, or neither. *WHRN* is indicated with an open star. Dashed lines mark zero specificity in each cell family. A complete list of values is available in Supplementary table 3a. (b) Heatmap showing single-cell RNA expression (nCPM) of *WHRN* and the three partners exhibiting significant endocrine specificity in (A), across the 10 endocrine cell types shown in Figure 2. Cell colour indicates log₁₀(nCPM+1) and the corresponding nCPM value for each gene/cell-type pair is displayed within each cell, also available in Supplementary table 3b.

To determine whether the endocrine specificity of *ADGRV1*, *PCDH15,* and *USH2A* reflects a broad elevation across endocrine cell types, or is driven by a single outlier, we compared their single-cell expression levels (nCPM), alongside those of *WHRN*, across all 10 endocrine cell types (Figure 3b). *ADGRV1* was broadly elevated (488-789 nCPM) across endocrine cells, with high expression in adrenal cortex cells and the five hormone-producing anterior pituitary cell types (gonadotrophs, thyrotrophs, somatotrophs, lactotrophs and corticotrophs), but comparatively low levels in adrenal medulla cells (21 nCPM), neuroendocrine cells (55 nCPM), pancreatic islet cells (90 nCPM) and Leydig cells (1.0 nCPM). *PCDH15* followed a narrower but still multi-cell-type pattern, with elevated expression in adrenal cortex cells (1,027 nCPM), lactotrophs (1,040 nCPM), adrenal medulla cells (511 nCPM) and thyrotrophs (443 nCPM), while remaining relatively low in the six remaining endocrine cell types (4.3-191 nCPM). In contrast, *USH2A* showed low expression in nine of the ten endocrine cell types (1.1-36 nCPM) with its high endocrine specificity rank driven primarily by a single cell type, adrenal medulla cells (223 nCPM), where expression was 6- to >100-fold higher than in any other endocrine cell type.

### Protein immunohistochemistry in cancerous tissues does not reveal a general overexpression but rather a trend in specific cases

To assess whirlin expression in cancerous tissues, IHC was performed across 20 common cancer types. Figure 4a reports IHC-based annotations of whirlin expression in the different cancer tissues, and Figure 4b displays representative IHC staining images.

**Figure 4:**
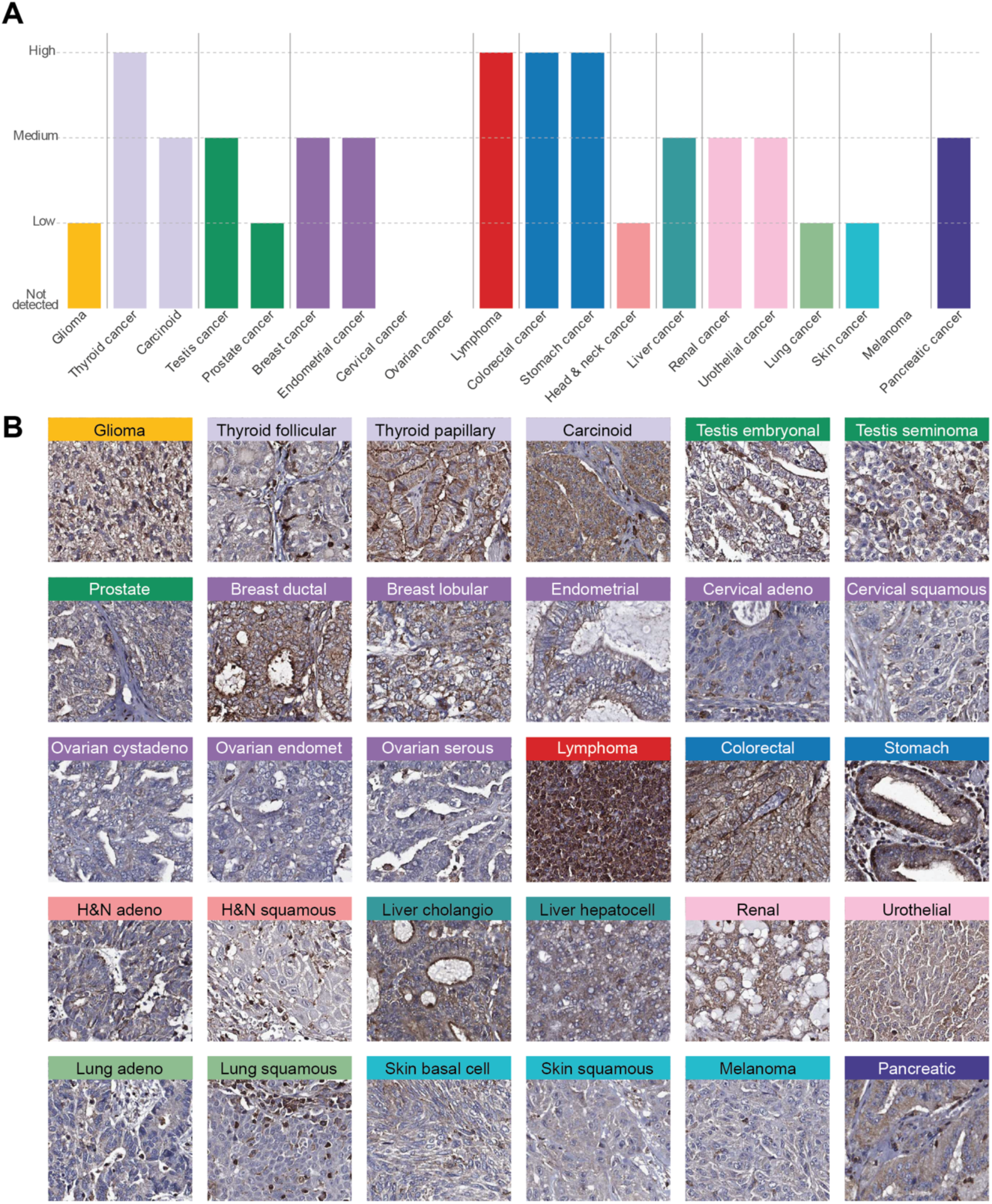
Whirlin protein expression across human cancer types. (a) IHC-based annotation scores from the HPA are displayed for 20 cancer types, representing the most frequently observed annotation level across patient samples (Not detected / Low / Medium / High). Detailed expression annotations for all cancer types and patient samples are provided in Supplementary Table 4. (b) Representative IHC images showing whirlin expression across the analysed cancer types.

Overall, whirlin was detected in a wide range of cancers, with expression levels generally comparable to those observed in the corresponding normal tissues without evidence of consistent overexpression. Although high expression was detected in individual cases, staining intensity often varied between patients within the same cancer type. In addition, it should be noted that several cancer types were represented by multiple histological subtypes.

The strongest expression was observed in lymphoma, where 10 out of 12 patients showed high staining levels. High expression was also detected in colorectal and stomach cancers, with high-to-medium staining observed in all 10 colorectal cancer samples and in 8 out of 12 stomach cancer samples. Similarly, high staining was detected in 2 out of 3 thyroid cancer samples. A detailed report of expression across all cancer types and individual patients is provided in Supplementary Table 4.

Medium-to-high staining was additionally detected in several adenocarcinomas, including liver, pancreatic, urothelial, renal, breast, and endometrial cancers. Representative IHC images illustrate prominent cytoplasmic staining within cancerous cells in these tissues. Lower levels were observed in glioma, prostate cancer, all squamous cell carcinomas, and melanoma where staining was weak or detected only in a minority of patients.

### Multiple human cell lines display endogenous WHRN expression

To identify in vitro models expressing endogenous *WHRN*, RNA expression levels were analysed across a large panel of human cell lines included in the HPA cell line resource (see Supplementary Table 5). This dataset includes data from a wide range of cancerous and non-cancerous cell lines of multiple tissue origins.

Overall, *WHRN* was detected across a broad spectrum of cell lines. The highest expression was observed in myeloma-derived cell lines, where *WHRN* RNA levels were markedly elevated compared to most other cell lines. Here, 15 of the 34 myeloma cell lines exceeded 65 nTPM, with the highest reaching 188 nTPM. High expression (> 60 nTPM) was also observed in two lymphoma cell lines (OCI-Ly10 and Pfeiffer), two skin cancer cell lines (LOX-IMVI and DEOC-1) and one rhabdoid cell line (BT-12). Moderately high expression (> 40 nTPM) was further detected in isolated cell lines from breast, lung and bone cancers. More broadly, epithelial cancer cell lines from different organs showed moderate *WHRN* expression, whereas cell lines derived from endocrine cancers and brain tumours tended to show low levels. Notably, several standard human cell lines expressed endogenous *WHRN* RNA, including RPMI-8226, HeLa, MCF-7, PC-3 and HEK293.

## DISCUSSION

Although scaffold Usher proteins such as whirlin are well characterised for their roles in hearing and vision, their expression and potential functions in other organ systems remain largely unexplored. In this study, we present the first comprehensive expression landscape of whirlin across most major human normal tissues and cancer types at single-cell level by integrating transcriptomics and antibody-based proteomics data. The data generated here was incorporated into version 25 of the HPA database, providing a publicly accessible resource for the broader research community, and the possibility to further explore the spatial localisation in high-resolution histological images.

Bulk RNA analyses and IHC showed that whirlin is detected in most human tissues, with particularly high levels observed in endocrine organs, reproductive tissues, and some regions of the brain and nervous system. Single-cell RNA analyses further defined this tissue-wide distribution by revealing that *WHRN* is strongly enriched in specific cell types rather than uniformly distributed across tissues.

We specifically observed high expression levels in ciliated cells and specialised epithelial cells across several organs. This distribution is notable because whirlin is known to localise in specialised actin-rich and ciliary structures in sensory cells, such as the stereocilia of inner ear hair cells and the connecting cilium in photoreceptors.[5] Genome-wide, *WHRN* ranked among the top 3.1% most cell-type-specific genes for ciliated cells, and a 500-gene module built from genes co-varying with *WHRN* across the atlas showed enrichment for cilium assembly, intraflagellar transport, basal-body anchoring, and BBSome-mediated cargo targeting. This pattern replicated in the independent compendium of WebCSEA, where ciliated-cell panels ranked among the most enriched overall. Together, these results indicate that whirlin’s association with ciliated cells is not confined to specific sensory cells, suggesting that whirlin may have a broader role in the organisation or maintenance of membrane-cytoskeletal structures associated with cellular protrusions or polarised epithelial functions. Consistent with this interpretation, enrichment was also observed in epithelial cells involved in secretion, absorption, or barrier functions, including glandular epithelial cells of digestive and reproductive tissues.

Another notable finding was the strong enrichment of *WHRN* in endocrine cells, including adrenal cortex cells and multiple hormone-producing pituitary cell types. Genome-wide specificity analysis placed *WHRN* among the top 3.4% of genes for endocrine-cell specificity. Importantly, this endocrine enrichment extended to several known binding partners of whirlin. To our knowledge, endocrine-cell specificity of *WHRN* and other Usher genes has not been previously reported. Although these observations remain preliminary, calling for further proteomics studies, the possible enrichment of a multi-domain scaffolding protein such as whirlin in these cells, together with its binding partners, raises the possibility that they may contribute to the spatial organisation of signalling or trafficking complexes involved in hormone secretion. This observation echoes a clinical report linking whirlin to Hashimoto’s thyroiditis.[15] In addition, our observation of high expression in thyroid cancer tissues raises the question of whether whirlin might be upregulated in some endocrine cancers. Further analyses on extended tissue samples containing all the specialised regions of these tissues, including the various functional layers of adrenal gland, would elucidate the potential role of whirlin and partners in these tissues.

Our analysis of cancer tissues points to expression levels generally mirroring those observed in corresponding normal tissues, suggesting that whirlin’s presence in tumours largely reflects the cellular origin of the cancer rather than a cancer-associated upregulation. Strong staining was however observed in several tumour types, particularly lymphoma and gastrointestinal cancers such as colorectal and stomach cancers. These observations align with an earlier study reporting the upregulation of several whirlin isoforms in colorectal tissues during colorectal cancer progression.[18] Subsequent pharmacogenomic analyses have also identified *WHRN* variants associated with differential responses to cetuximab-based therapy in colorectal cancer,[24] while more recent transcriptomic studies have reported dysregulated *WHRN* expression in gastric cancer.[25] It should be noted that the investigated tumours in our study only contained up to 12 individual samples per tumour type. Considering the heterogeneity of tumours and various subtypes, this number is too small to be able to draw a biological conclusion about the expression pattern across tumours, and highlights the importance of future studies on larger patient cohorts.

The identification of multiple human cell lines expressing endogenous *WHRN* provides practical experimental models for future studies. High expression levels were detected in hematopoietic-derived cell lines, particularly myeloma and lymphoma cells. Most importantly, *WHRN* was detected in widely used laboratory cell lines such as HEK293, HeLa, and MCF-7, providing accessible systems for investigating the molecular functions of whirlin. A complete list of *WHRN* expression levels in 154 cell lines can be found on the cell line panel of the protein database for detailed analyses for the scientific community.

Together, this study expands the current view of whirlin biology by demonstrating that it is broadly expressed across human tissues and enriched in specific cell types beyond sensory systems. The expression landscape presented here argues that whirlin could act as a ubiquitous membrane scaffold whose full biological role has been underestimated. This work establishes a foundation for future investigations aimed at understanding the broader physiological and pathological roles of this Usher syndrome protein and its binding partners.

## MATERIALS AND METHODS

### Data sources

Normalised *WHRN* RNA expression data for tissues,[26] brain,[27] single-cell,[19] and cell line resources[28] were retrieved from the Human Protein Atlas (HPA) database (https://v25.proteinatlas.org) and are all gathered in the Supplementary Tables 1-5.

Whirlin protein expression levels based on IHC correspond to staining intensity scores through manual annotation, as described below.

### Human tissues and sample preparation

Human tissue samples used for protein expression analysis in the HPA datasets were collected and handled in accordance with Swedish laws and regulations. Tissues were obtained from the Clinical Pathology Department at Uppsala University Hospital, Sweden, and collected through the Uppsala Biobank organisation. All samples were anonymised in accordance with approvals and advisory reports from the Uppsala Ethical Review Board (Ref. #2002-577, 2005-388, 2007-159, 2011-473). Informed consent was obtained from all study participants.

For immunohistochemical analysis, formalin-fixed, paraffin-embedded (FFPE) tissue blocks from pathology archives were selected based on normal histology, as evaluated on hematoxylin-eosin-stained sections. Representative 1-mm-diameter cores were sampled from FFPE blocks and assembled into tissue microarrays (TMAs) containing normal or cancer tissue samples.

Three TMAs were analysed for the antibody used in this study: one containing 45 normal tissue types, another including 20 cancer types, and a third specialised TMA with exclusively retina tissue. Each normal tissue was represented by three cores, except for retina which was represented by two cores, and cancer tissues were represented by two cores per sample, with up to 12 patients included for each cancer type.

### Immunohistochemistry

Immunohistochemical staining and high-resolution digitisation of TMA slides were performed as previously described[29]. TMA blocks were sectioned at 4 µm using a waterfall microtome (Microm H355S, Thermo Fisher Scientific, Fremont, CA), mounted on SuperFrost Plus slides (Thermo Fisher Scientific), dried overnight at room temperature, and baked at 50 °C for at least 12 h.

Automated staining was carried out using a Lab Vision Autostainer 480S module (Thermo Fisher Scientific, Fremont, CA) following previously described protocol.[29] A polyclonal rabbit IgG antibody against human whirlin (CAB080582 / 25881-1-AP, Thermo Fisher Scientific, Fremont, CA) was used as the primary antibody. Antibody dilution and staining conditions were optimised according to the HPA pipeline for antibody validation,[30] using two test TMAs containing in total 20 different normal tissues. A dilution of 1:800 was selected.

Slides were digitised with a ScanScope AT2 system (Leica Aperio, Vista, CA) using a 20x objective. All tissue samples were manually annotated by RS, BK and CL, following the standardised HPA workflow.[20] Staining intensity was used as the main scoring parameter on a four-level scale: 0 = not detected, 1 = low, 2 = medium, and 3 = high.

### Cell-type classification

*WHRN* normalised counts per million (nCPM) across 154 single-cell types, compiled by the HPA from 34 published single-cell RNA-seq datasets, were retrieved from the HPA single-cell dataset (Supplementary Table 2). A threshold of 50 nCPM was used throughout to define cell types with substantial *WHRN* expression (Figure 2a). Each cell type was independently assigned to one of five protrusion classes describing its dominant apical surface specialisation: motile cilia, flagellum, sensory cilium, microvilli/brush border, or no specialised protrusion, curated from HPA cell-type annotations and the literature (Figure 2b; full classification in Supplementary Table 2). For the gene-level specificity analyses (Figures 2c and 3a), cell families were defined as follows: cell types bearing motile cilia, a flagellum, or a sensory cilium were grouped into a single ciliated/flagellated family (n = 10 ciliated cell types), and for the endocrine family, ten independently curated endocrine cell types were retained (adrenal cortex cells, adrenal medulla cells, gonadotrophs, thyrotrophs, somatotrophs, lactotrophs, corticotrophs, neuroendocrine cells, pancreatic islet cells, and Leydig cells); the remaining 134 cell types formed the comparison group for both tests.

### Gene-level cell-type specificity ranking

Genome-wide cell-type specificity was assessed using a one-vs-rest differential expression test. Of the 154 cell types, genes expressed at nCPM ≥ 1 in at least 5 cell types were retained (17,984 genes). For each gene and each cell family (ciliated or endocrine), expression was natural-log transformed (log1p(nCPM)) and the median across the 10 family cell types was compared with the median across the remaining 144 using a one-sided Mann-Whitney U test; the difference between these two medians is reported as the specificity value. P-values were corrected for multiple testing across all 17,984 genes using the Benjamini-Hochberg procedure. Genes were ranked by specificity value within each family (rank 1 = highest), and *WHRN’s* percentile rank was determined from this genome-wide distribution. The same test, applied to the same 17,984-gene background, was used to rank each of 13 known *WHRN* binding partners (listed in Supplementary Table 6) for both ciliary and endocrine specificity.

### Co-expression module and pathway enrichment

Genes with expression patterns similar to *WHRN* across the 154 cell types were identified by pairwise Spearman correlation; of 17,367 genes with usable expression data, 3,002 showed significant co-variation with *WHRN* (q < 0.01, Benjamini-Hochberg corrected). A co-expression module was defined as the 500 genes with the strongest positive correlation (Spearman ρ ≥ 0.37). Enrichment of cilium-related Gene Ontology annotations within this module was assessed using Fisher’s exact test against a background of genes matched for both mean expression and detection breadth across the 154 cell types, rather than an unmatched random sample. The 500-gene module was submitted to the Reactome Pathway Analysis Service (reactome.org/AnalysisService) for over-representation analysis against curated Reactome pathway gene sets, using default hypergeometric testing with Benjamini-Hochberg FDR correction.

### WebCSEA

The 500-gene co-expression module was tested for independent replication using WebCSEA (https://bioinfo.uth.edu/webcsea/),[23] a cell-type-specific expression enrichment tool that benchmarks a submitted gene list against 1,355 cell-type panels, each an independent (cell type × originating study) pair, drawn from 111 published single-cell RNA-seq datasets, using its default combined-P statistic per panel. Panels were classified into ciliated, endocrine, or retinal families by matching panel and cell-type annotations against curated keyword lists (e.g. "cilia" for ciliated panels; organ-specific terms for endocrine panels). Bonferroni correction was applied across all 1,355 panels tested, and family-level enrichment was assessed by comparing each family’s panel ranks against all remaining panels using a one-sided Mann-Whitney U test.

### Data visualisation

All figures were generated in R (version 4.4.2) using the ggplot2 package (version 4.0.1). Tissues and cell types were coloured according to HPA classification scheme. Gene-level specificity ranking, co-expression analysis, GO/Reactome enrichment, and WebCSEA replication were performed in Python (pandas 2.3.3, numpy 2.4.6, scipy 1.17.1, statsmodels, requests), in addition to the R/ggplot2 pipeline used for figure rendering.

### Data availability

All datasets analysed in this study are publicly available through the HPA database (https://v25.proteinatlas.org) and WebCSEA (https://bioinfo.uth.edu/webcsea/). IHC staining images for all normal and cancer tissues are available in the HPA and have been included since version 25 of the database.

## ASSOCIATED CONTENT

### Supporting Information

The following files are available free of charge: Supplementary Tables 1-5

- Supplementary Table 1a: *WHRN* bulk RNA expression from the consensus HPA dataset.
- Supplementary Table 1b: Whirlin protein expression annotation from IHC in normal human tissues.
- Supplementary Table 2: *WHRN* single-cell RNA expression from the HPA single-cell dataset, including each cell type’s protrusion class.
- Supplementary Table 3a: Ciliary and endocrine cell-type specificity rank, specificity value, and FDR for *WHRN* and its 13 known binding partners.
- Supplementary Table 3b: Single-cell RNA expression of *WHRN, ADGRV1, PCDH15,* and *USH2A* across the 10 endocrine cell types.
- Supplementary Table 4: Whirlin protein expression annotation from IHC in cancerous tissues.
- Supplementary Table 5: *WHRN* cell lines RNA expression from the HPA cell line dataset.
- Supplementary Table 6: Whirlin Partners List

## AUTHOR INFORMATION

### Author Contributions

Conceptualization: C.G.; Data curation: A.D & C.G.; Formal analysis: A.D, L.M, C.L, C.G.; Methodology: L.M.; Writing - original draft: C.G.; Writing - review & editing: the manuscript was reviewed and edited through contributions of all authors. All authors have given approval for the final version of the manuscript.

## ACKNOWLEDGMENT

The authors would like to thank all the staff of the Human Protein Atlas for their work specifically Rutger Schutten and Borbala Katona, and the patients giving tissues for research.

## ABBREVIATIONS

BBSome: Bardet-Biedl Syndrome protein complex
CSEA: Cell-type Specific Enrichment Analysis
DFNB31: Deafness Autosomal Recessive 31
FDR: False Discovery Rate
FFPE: Formalin-Fixed Paraffin-Embedded
GO: Gene Ontology
GTEx: Genotype-Tissue Expression
HHD: Harmonin Homology Domain
HPA: Human Protein Atlas
IHC: Immunohistochemistry
IWGAV: International Working Group for Antibody Validation
nCPM: Normalized Counts Per Million
nTPM: Normalized Transcripts Per Million
scRNA: single-cell RNA
TMA: Tissue microarray
USH2: Usher Syndrome Type 2
WebCSEA: Web-based Cell-type-Specific Enrichment Analysis.

